# Regulation of Lysosome-Associated Membrane Protein 3 (LAMP3) in Lung Epithelial Cells by Coronaviruses (SARS-CoV-1/2) and Type I Interferon Signaling

**DOI:** 10.1101/2021.04.28.441840

**Authors:** Chilakamarti V. Ramana

## Abstract

Severe acute respiratory syndrome coronavirus-2 (SARS-CoV-2) infection is a major risk factor for mortality and morbidity in critical care hospitals around the world. Lung epithelial type II cells play a major role in several physiological processes, including recognition and clearance of respiratory viruses as well as repair of lung injury in response to environmental toxicants. Gene expression profiling of lung epithelial type II-specific genes led to the identification of lysosomal-associated membrane protein 3 (LAMP3). Intracellular locations of LAMP3 include plasma membrane, endosomes, and lysosomes. These intracellular organelles are involved in vesicular transport and facilitate viral entry and release of the viral RNA into the host cell cytoplasm. In this study, regulation of LAMP3 expression in human lung epithelial cells by several respiratory viruses and type I interferon signaling was investigated. Coronaviruses including SARS-CoV-1 and SARS-CoV-2 significantly induced LAMP3 expression in lung epithelial cells within 24 hours after infection that required the presence of ACE2 viral entry receptor. Time-course experiments revealed that the induced expression of LAMP3 by SARS-CoV-2 was correlated with the induced expression of interferon-beta1 (IFNB1) and signal transducers and activator of transcription 1 (STAT1) mRNA levels. LAMP3 was also induced by direct IFN-beta treatment or by infection with influenza virus lacking the non-structural protein1(NS1) in NHBE bronchial epithelial cells. LAMP3 expression was induced in human lung epithelial cells by several respiratory viruses, including respiratory syncytial virus (RSV) and the human parainfluenza virus 3 (HPIV3). Location in lysosomes and endosomes as well as induction by respiratory viruses and type I Interferon suggests that LAMP3 may have an important role in inter-organellar regulation of innate immunity and a potential target for therapeutic modulation in health and disease. Furthermore, bioinformatics revealed that a subset of lung type II cell genes were differentially regulated in the lungs of COVID-19 patients.

## 1. Introduction

The lung is a complex organ composed of many cell types including epithelial, endothelial, fibroblast, and smooth muscle origin (1,2). Alveolar epithelium provides a barrier function to invading respiratory viruses and consists of two morphologically and functionally distinct type I and type II cells (3–5). Human lung type II cells are highly susceptible to SARS-CoV-2 infection as they co-express Angiotensin-converting enzyme2 (ACE2) required for virus entry and TMPRSS2 protease involved in the cleavage of the spike protein of the virus (6). Lung type II cells provide a protective function to the lung by detoxification of air and chemical pollutants that reach the lungs by inhalation or via circulation (7). In addition, lung type II cells secrete a variety of anti-inflammatory and antimicrobial substances into the alveolar fluid such as surfactant proteins, lysozyme, lipocalin2, and reduced glutathione (8). Lung type II cells can undergo cell proliferation and differentiate into type I cells in response to lung injury (9) Lung type II cells are characterized by cell-specific expression of surfactant protein C (SP-C) that is a major source of surfactant proteins and reduces surface tension (10). The sequence elements in the SP-C gene promoter mediate tissue-specific expression of heterologous genes such as influenza hemagglutinin (HA) antigen in lung type II cells in transgenic mice (11). This approach was used to dissect the role of cytokines and transcription factors involved in CD8^+^ T cell-mediated lung injury (12,13). Considerable progress has been made in understanding the role of type II cells in lung function and diseases. The development of type I and type II cell-specific quantitation methods facilitated investigating the role of alveolar epithelium in lung injury and repair (14). Gene expression profiling studies revealed a large number of lung type II cell-specific genes encoding several proteins located in membrane compartments such as ion channels, transporters, and vesicular proteins like lysosome-associated membrane protein 3 or LAMP3 (2,6,15). In this study, regulation of LAMP3 by respiratory viruses including Coronaviruses SARS-CoV-1 and SARS-CoV-2 as well as type I Interferon (IFN-α/β) signaling was investigated in human lung epithelial cells and in the lung tissue samples of healthy and COVID-19 patients.

## 2 Materials and Methods

### 2.1 Gene expression datasets and Bioinformatics

Mouse lung type II- specific gene data by single-cell RNA sequencing was described previously (6). The data was downloaded from LUNGGENS website at https://research.cchmc.org/pbge/lunggens/. Gene expression in mouse lung type II cells by microarrays was reported previously (15). Gene expression datasets representing human lung cell lines infected with respiratory viruses and from lung biopsies of healthy and COVID-19 patients were reported previously (16,17). Gene expression resources from Immgen RNA seq SKYLINE were used (http://rstats.immgen.org/Skyline_COVID-19/skyline.html). Outliers of expression were not included in the data analysis. Supplementary data was downloaded from the Journal publisher website and Geo datasets were analyzed with the GeoR2R method (NCBI). Cluster analysis was performed using gene expression software tools (18). Protein-protein interactions were visualized in BIOGRID or the Search Tool for the Retrieval of Interacting Genes/Proteins (STRING) databases (19,20). Gene ontology (GO) and transcription factor analysis were done in Metascape and TRANSFAC databases, respectively (21,22). Interferon-related data was retrieved from WWW.Interferome.org. Gene-specific information was retrieved from standard bioinformatics websites such as Mouse Genome Informatics (MGI), Human Protein Atlas, GTEx database, TCGA cancer database, and GeneCards.

## 3. Results and Discussion

### 3.1 Profiling Lung epithelial type II cell-specific transcriptome

Microarrays and single-cell RNA sequencing provided novel insights into the gene expression profile in mouse primary lung type II cells. Functional organization of the lung type II transcriptome revealed that many genes were involved in metabolism, membrane transport, detoxification, immune responses and cell growth (6,15). Furthermore, several genes were exclusively expressed in lung type II cells. Gene ontogeny (GO) term analysis revealed that these genes were involved in lipid, carbohydrate, and cholesterol metabolism (Figure 1A). Cell-specific expression of some of these genes in Clara, Ciliated, lung type II (AT2), lung type I (AT1) cells were shown (Figure 1B). Expression analysis in widely used human lung epithelial cell lines revealed that surfactant genes were not expressed in these cell lines. However, membrane proteins like LAMP3 and solute carrier family 34 member 2 (SLC34A2) as well as regulators of lipid and carbohydrate metabolism like lipocalin 2 (LCN2), lysozyme (LYZ) were expressed at significant levels in the Calu3 cell line than in A549 or NHBE cell lines (Figure 1C). Therefore, Calu3 represents a useful cell culture system to study the regulation of these genes under a variety of conditions.

**Figure 1.**
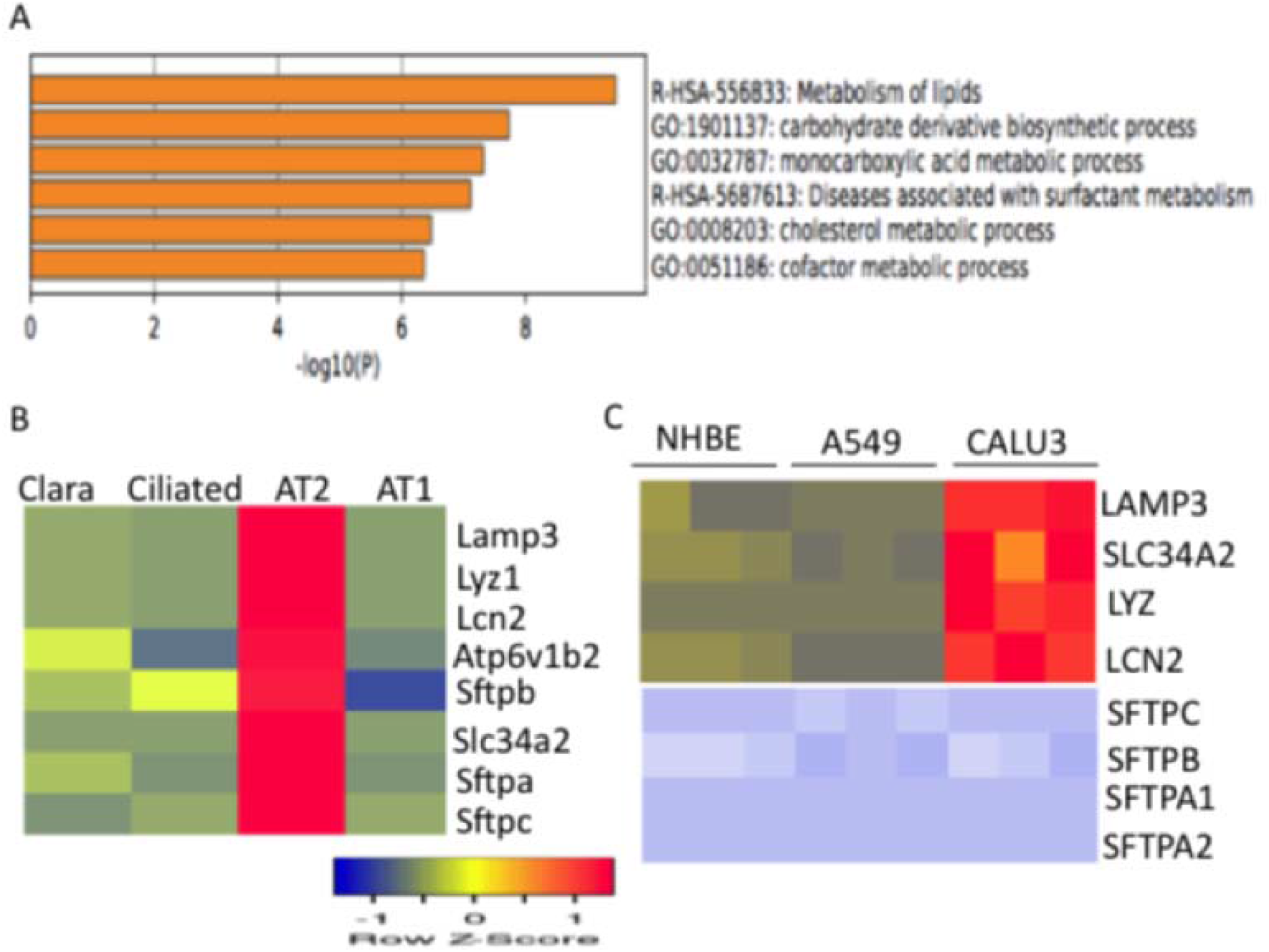
Annotation of gene ontology (GO) terms and biological functions associated with mouse lung type II cell-specific transcriptome and regulation of gene expression in different cell types. (A) Biological functions and gene ontology (GO) terms associated with mouse lung type II cell-specific transcriptome were ranked by significance (B) Expression levels of lung type II cell-specific genes in Clara, Ciliated, AT2, and AT1 cells (C) Expression levels of lung type II epithelial-specific genes in A549, NHBE, and Calu3 human cell lines. Gene expression profiles were retrieved from gene array datasets. Cluster analysis was performed with the Heatmapper software.

### 3.2 Tissue distribution and intracellular location of LAMP3

LAMP3 was originally identified as a human lung-specific gene located on chromosome 3 with homology to lysosomal membrane proteins LAMP1 and LAMP2 and expressed in several cancer tissues (23). Tissue distribution analysis in a large set of human tissues in the GTEx database (Broad Institute, Cambridge, MA) confirmed that LAMP3 was highly expressed in the human lung and in Epstein-Barr virus (EBV)-transformed B lymphocytes (Figure 2A). Tissue distribution of LAMP3 protein in the Human Protein Atlas database showed high levels of expression in the lung and moderate levels of expression in the lymph nodes and tonsils (Figure 2B) Annotation in TCGA cancer database indicated that LAMP3 was detected in many cancers and exhibited low tissue specificity. The prognosis of cancer patients as represented by Kaplan-Meier survival plots revealed that LAMP3 gene expression as a marker for renal cancer is unfavorable and for ovarian cancer as favorable (data not shown).

**Figure 2.**
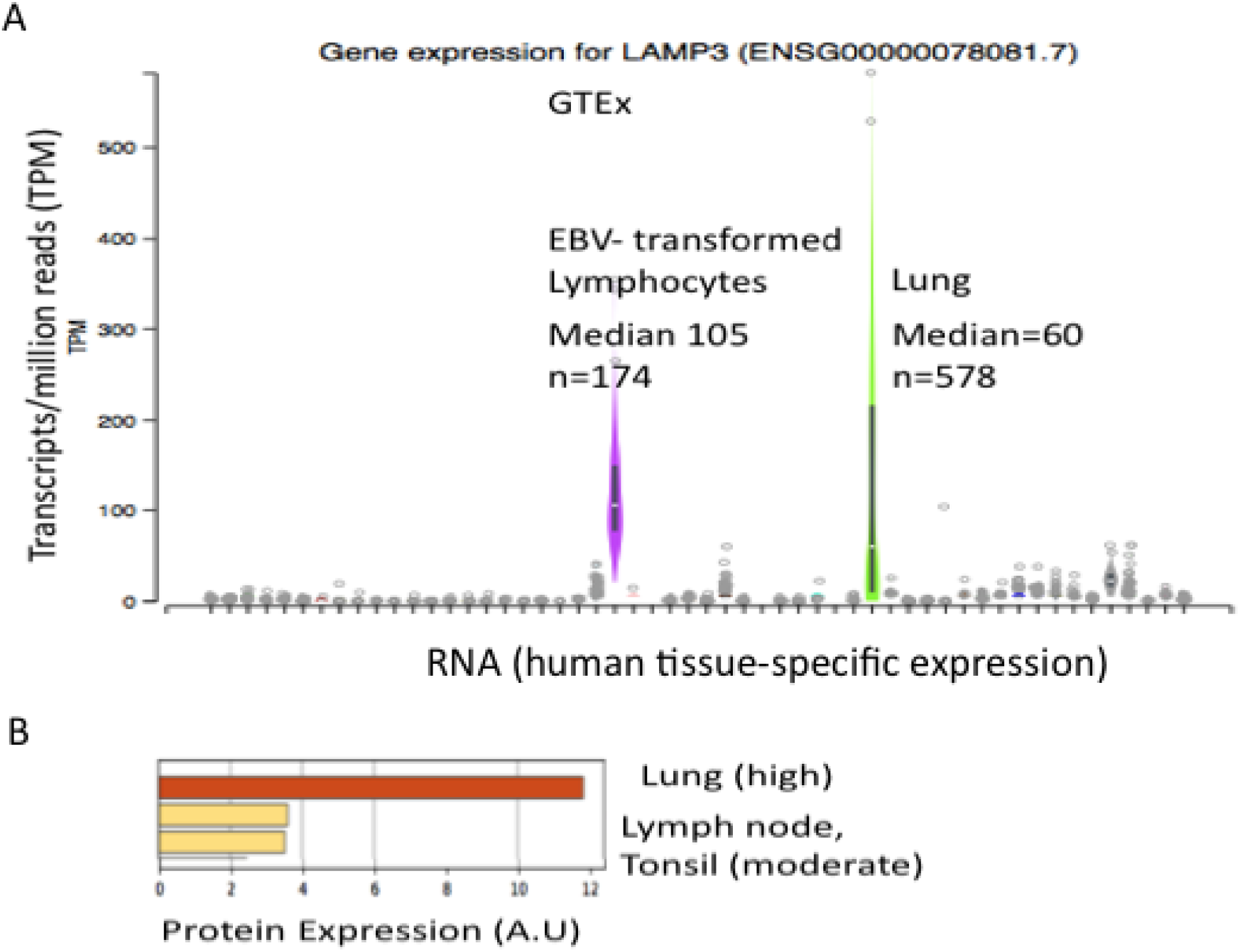
Expression of LAMP3 mRNA levels in human tissues. (A) LAMP3 mRNA expression levels in human tissues were represented in the GTEx database. Transcript levels in million reads (TPM) were shown (B) LAMP3 protein levels in human tissues were represented in the Human Protein Atlas database.

Vesicles are organelles enclosed by a lipid bilayer in the cell and vesicular transport represents a major pathway of intracellular transport. Lipid and protein components of surfactant are secreted via fusion of the vesicles with the plasma membrane by a process known as exocytosis and part of the used surfactant is recycled via endocytosis (24). The specialized vesicles of lung type II cells are known as lamellar bodies and are coated with a variety of proteins including clathrin and adaptor protein (AP) complexes and regulated by proteins of the Ras/Rab family, ADP-ribosylation factor (ARF) family, and their effectors by post-translational modifications such as prenylation, palmitoylation, and ADP-ribosylation (25). SARS-CoV-2 binds to ACE2 and after proteolytic processing enters the cell via endosomes and lysosomes and the viral RNA is released into the cytoplasm for replication (26). These steps were schematically illustrated (Figure 3A). The intracellular location of a protein provides insights into the biological function of the protein. Annotation of intracellular distribution in various databases indicated that LAMP3 was located in various cell compartments such as vesicles, lysosomes, endosomes, plasma membrane, and lamellar bodies (Figure 3B).

**Figure 3.**
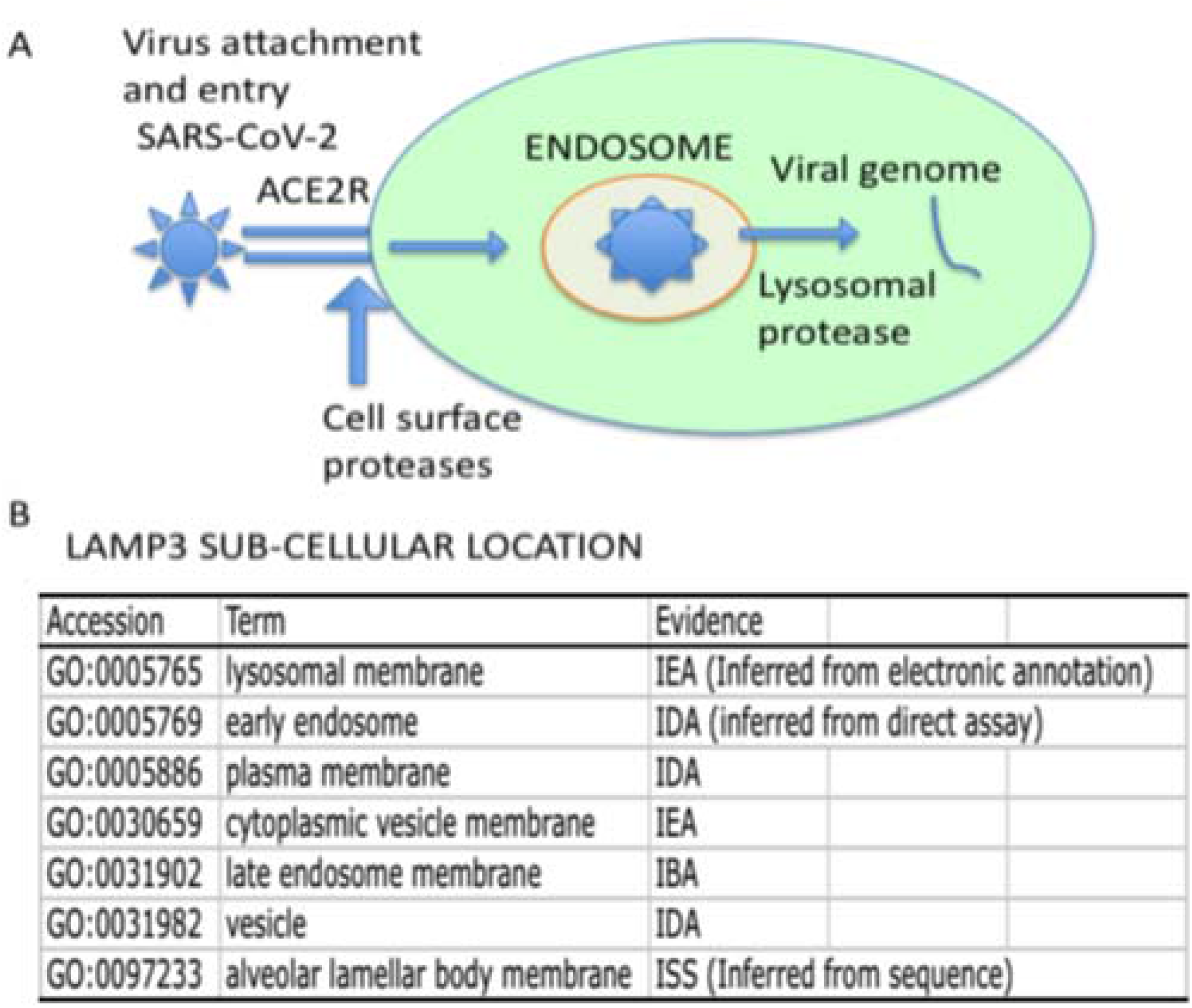
Intracellular location of LAMP3 protein. (A) Schematic outline of SARS-CoV-2 entry into the human lung epithelial cells. Important steps include binding of SARS-CoV-2 to angiotensin-converting enzyme-2 (ACE2) receptor, cleavage of virus spike protein by cell-surface proteases, vesicular transport of the virus through endosomes and lysosomes. Viral RNA is released into the cytoplasm (B) Intracellular location of LAMP3 protein were represented in the mouse genome informatics (MGI) and the human protein atlas databases.

### 3.3 Regulation of LAMP3 expression by pathogenic viruses and bacteria

Viruses and bacteria use endocytosis and vesicular trafficking to enter the host cell and release the nucleic acid into the cytoplasm. There is considerable evidence that LAMP3 plays a major role in the pathogenic responses to viruses and intracellular bacteria. Hepatitis C virus (HCV) infection as well as Interferon-α treatment induced LAMP3 expression during hepatic differentiation of pluripotent stem cells (27). Influenza A virus (IAV) infection induced LAMP3 expression in A549 human lung epithelial cells. Knock-down of LAMP3 expression by Si-RNA decreased viral nucleoprotein (NP) levels and attenuated viral replication and titers (28). Differential expression of LAMP3 in high and low responders to smallpox virus infection was reported (29). LAMP3 supports furin-mediated cleavage of the viral fusion protein of mumps virus in HEK 293 cells (30). *Mycobacterium tuberculosis* (Mtb) represents an intracellular mycobacterium and the causative agent of tuberculosis. Mtb was located in LAMP3-positive phagolysosomes of infected human macrophages (31). LAMP3 promotes the intracellular proliferation of *Salmonella typhimurium* (32). A genome-wide association study suggested LAMP3 is one of the risk loci for cutaneous leishmaniasis in Brazil (33).

### 3.4 Regulation of LAMP3 expression by coronaviruses SARS-CoV-1 and SARS-CoV-2

Coronaviruses and influenza viruses are the major respiratory viruses in humans with a potential pandemic threat and responsible for millions of deaths worldwide (34,35). Significant lung injury frequently accompanies respiratory virus infection, which is mediated by both the direct effects of the virus as well as a result of the host immune responses (36). The highly pathogenic SARS-CoV-2-induced gene expression signature in the human lungs was characterized by enhanced expression of Interferons and proinflammatory cytokines resulting in a dramatic increase in chemokines and inflammatory cell influx (37,38). Recognition of viral infection by the cell pathogen-associated molecular pattern (PAMP) receptors results in the phosphorylation and activation of IRF3 and induction of interferon-beta (IFNB1) gene transcription and secretion of type 1 interferon. The secreted or released interferon binds IFN α/β receptor on the surface of infected and neighboring cells to activate the Jak-Stat pathway to induce interferon-stimulated genes (39,40). Gene expression data in human lung epithelial cell lines such as Calu-3, A549, and NHBE1 in response to coronaviruses were reported (16,17). Interrogation of Microarray datasets revealed that LAMP3 was induced by SARS-CoV-1 and SARS CoV-2 infection but not by mock-infection in Calu-3 cells. This induction was observed at 12 and 24 hours after infection (Figure 4A and 4B). Interferon-beta (IFNB1) and transcription factors STAT1, STAT2, and IRF9 were also induced at mRNA levels by SARS-CoV-1 and SARS CoV-2 infection in Calu-3 cells in a time-dependent manner (Figure 5A and 5B). However, SARS-CoV-2 was more potent than SARS-CoV-1 in the induction of IFN-beta and ISGF3 components at mRNA levels in Calu3 cells. These results suggest that the production of type I interferon and autocrine activation of the Jak-Stat pathway plays a major role in the induction of interferon-stimulated genes. In contrast, SARS-CoV-2 infection of A549 failed to induce LAMP3 expression (Figure 6A). Previous studies have shown that ACE2 functions as the virus entry receptor and facilitates the entry of SARS-CoV1 and SARS-CoV-2 into lung epithelial cells (41). Expression of ACE2 in A549 cells rescued SARS-CoV-2-mediated LAMP3 induction and was inhibited by Ruxolitinib (Figure 6A). Furthermore, ACE2R expression dramatically increased the expression of IFNB1, STAT1, STAT2, and IRF9 mRNA after SARS-CoV-2 infection of A549 cells expressing ACE2 (Figure 6B). These results suggest that LAMP3 expression was dependent on IFNB1 production by SARS-CoV-2 in A549 cells and required the expression of SARS-CoV-2 entry receptor ACE2.

**Figure 4.**
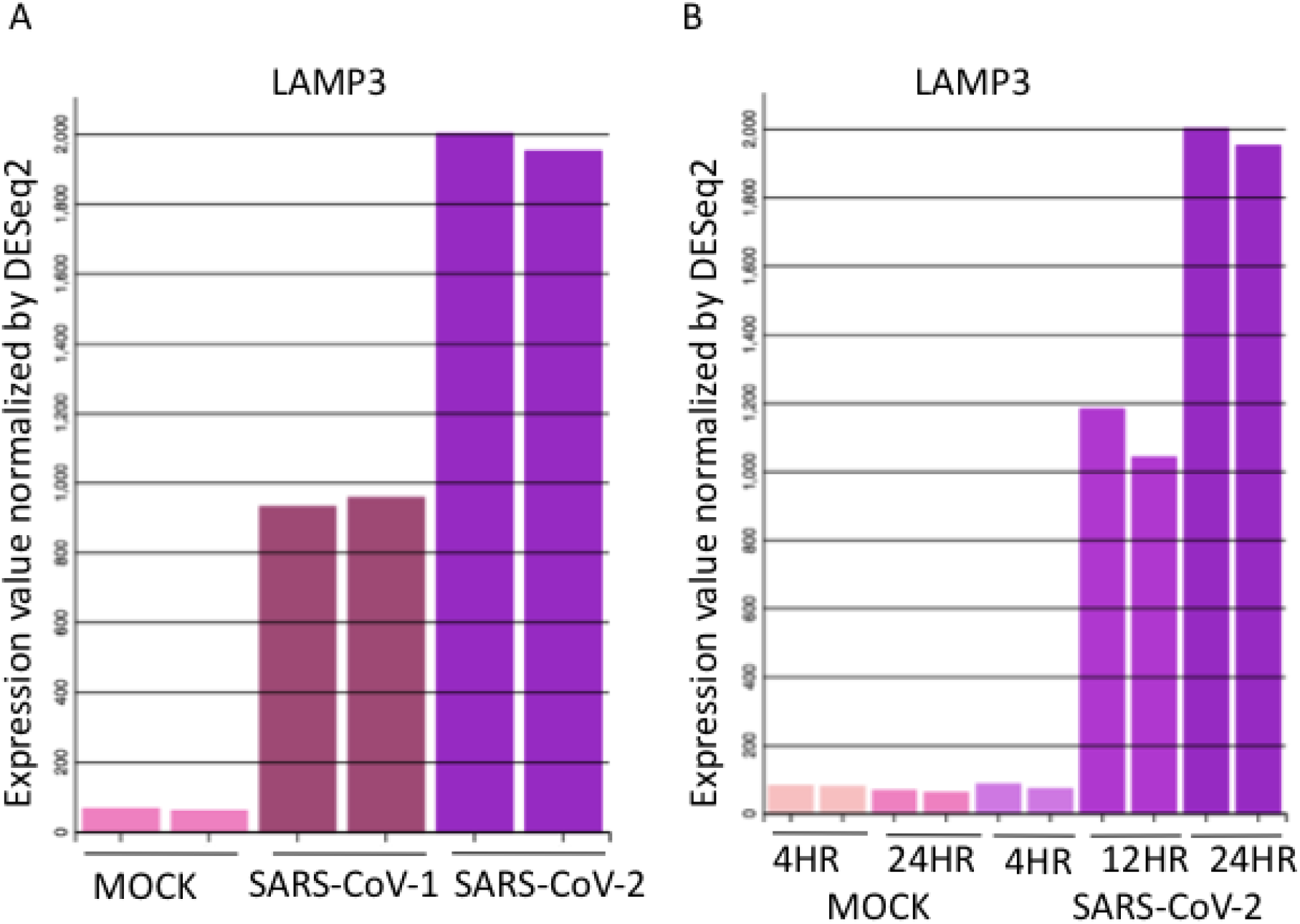
Regulation of LAMP3 mRNA levels by Coronaviruses in Calu-3 cells. (A) Calu-3 cells were mock-infected or infected with the virus (SARS-CoV-1 or SARS-CoV-2) for 24 hours. (B) Time-course of LAMP3 induction by SARS CoV-2 in Calu-3 cells. Mock-infection for 4 or 24 hours was used as controls. Data from two samples for each condition were shown. LAMP3 mRNA expression levels were normalized by DEseq2.

**Figure 5.**
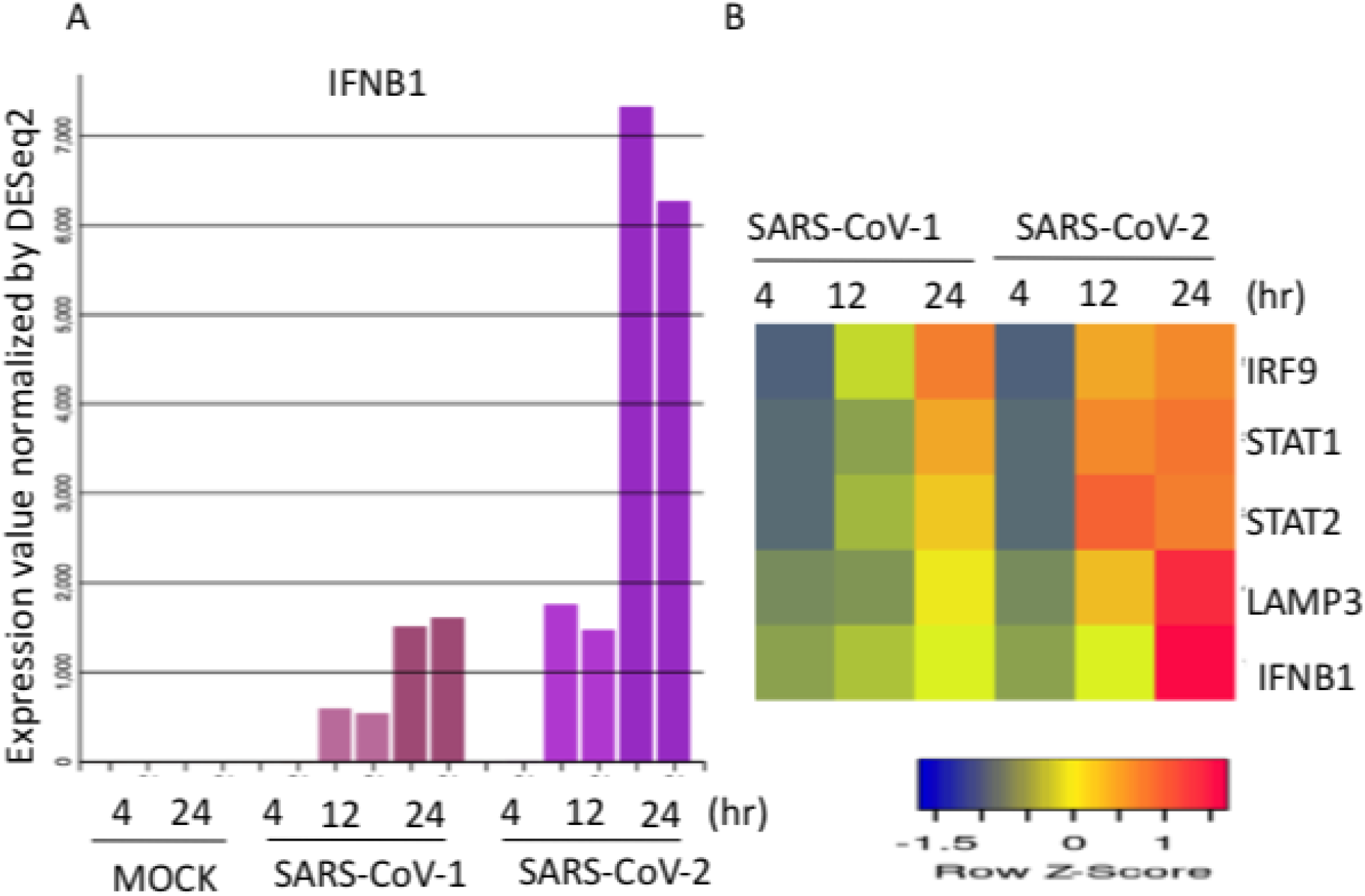
Regulation of IFN-β and ISGF3 (STAT1, STAT2, and IRF9) mRNA levels by SARS-CoV-1 and SARS CoV-2 in Calu-3 cells. (A) Calu-3 cells were mock-infected or infected with the virus (SARS-CoV-1 or SARS-CoV-2) for 4,12 or 24 hours. IFN-β mRNA expression levels were normalized by DEseq2. Data from two samples for each condition was shown. (B) Calu-3 cells were mock-infected or infected with the virus (SARS-CoV-1 or SARS-CoV-2) for 4-24 hours. The expression levels of LAMP3, IFNB1, and ISGF3 mRNA were normalized by DEseq2 method. Data from an average of two samples for each condition was shown.

**Figure 6.**
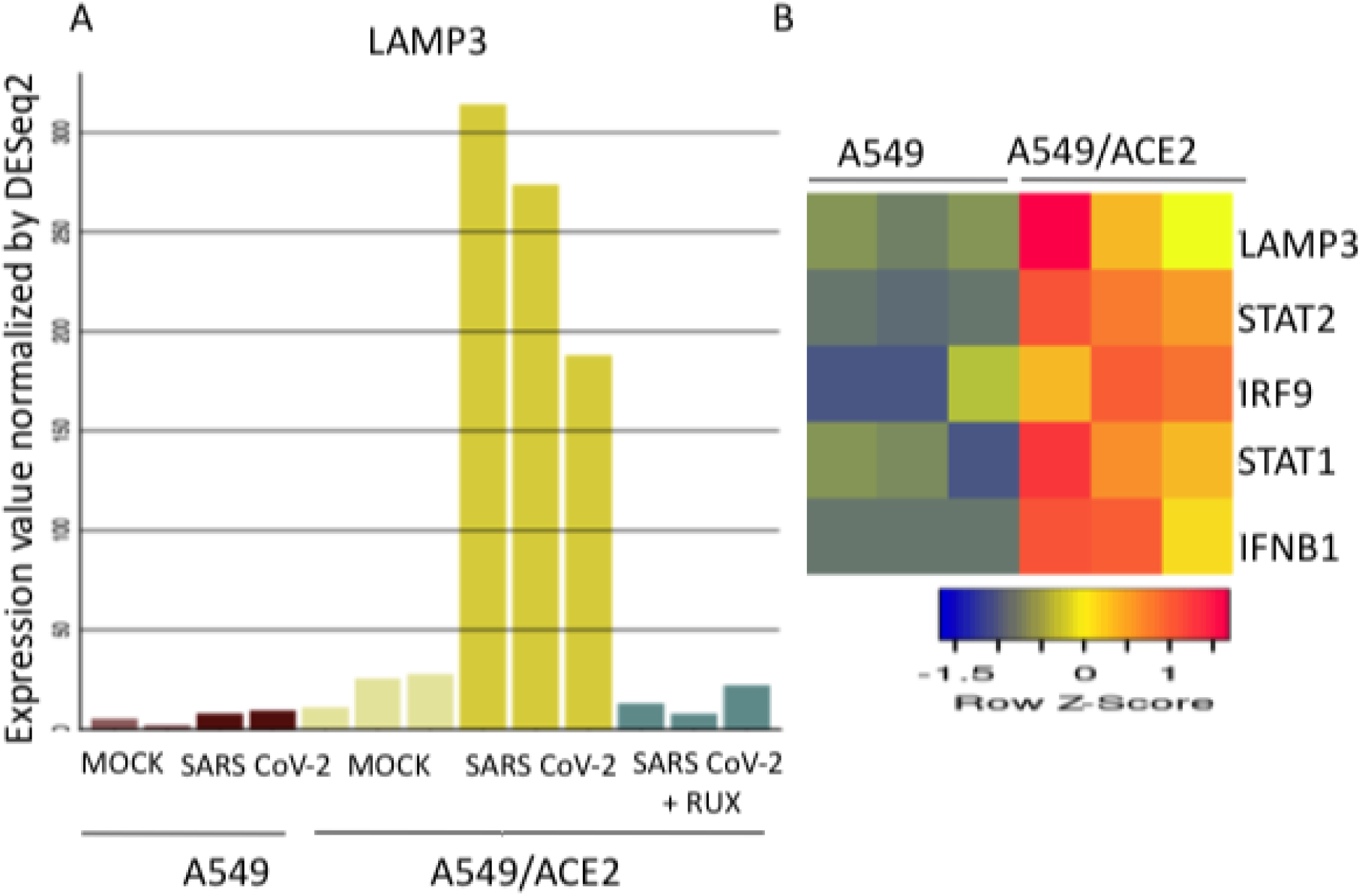
Regulation of LAMP3, IFNB1, and ISGF3 complex mRNA levels by SARS-CoV-2 in A549 or A549 cells expressing ACE2 receptors. (A) A549 or A549 cells expressing ACE2 receptors were mock-infected or infected with SARS-CoV-2 virus treated without or with Ruxolitinib (RUX) inhibitor for 24 hours. LAMP3 mRNA data from two or three samples for each condition were shown. (B) The expression levels of LAMP3, IFNB1, and ISGF3 mRNA were normalized by DEseq2 method. Data from three samples for each condition were shown.

### 3.5 Regulation of LAMP3 expression by Influenza A (IAV), Respiratory syncytial virus (RSV), and human parainfluenza virus (HPIV3)

Influenza A virus A (IAV) regulation of LAMP3 expression in NHBE bronchial epithelial cells was dependent on viral non-structural protein or NS1. wild-type IAV failed to induce LAMP3 expression in NHBE cells. In contrast, virus infection with a deleted NS1 resulted in the induction of LAMP3 expression (Figure 7A). Viral NS1 expression blocked the expression of IFNB1, STAT1, LAMP3, and interferon-stimulated gene expression (Figure 7B). NS1 inhibited the retinoic acid-inducible protein 1 (RIG-I) mediated induction of type 1 interferon (42). Furthermore, NS1 expression was associated with low activation of Stat1 and attenuation of interferon-stimulated gene expression (43). Respiratory syncytial virus (RSV) infection occurs mainly in children. In contrast, the human parainfluenza virus (HPIV3) can cause both upper and lower respiratory tract infections in children, immunocompromised adults, and the elderly (44). Despite these differences, both respiratory viruses induced LAMP3 expression in A549 cells (Figure 7C). These studies suggest that multiple factors such as virus strain, virus-encoded factors (such as NS1), cell type play an important role in viral pathogenesis.

**Figure 7.**
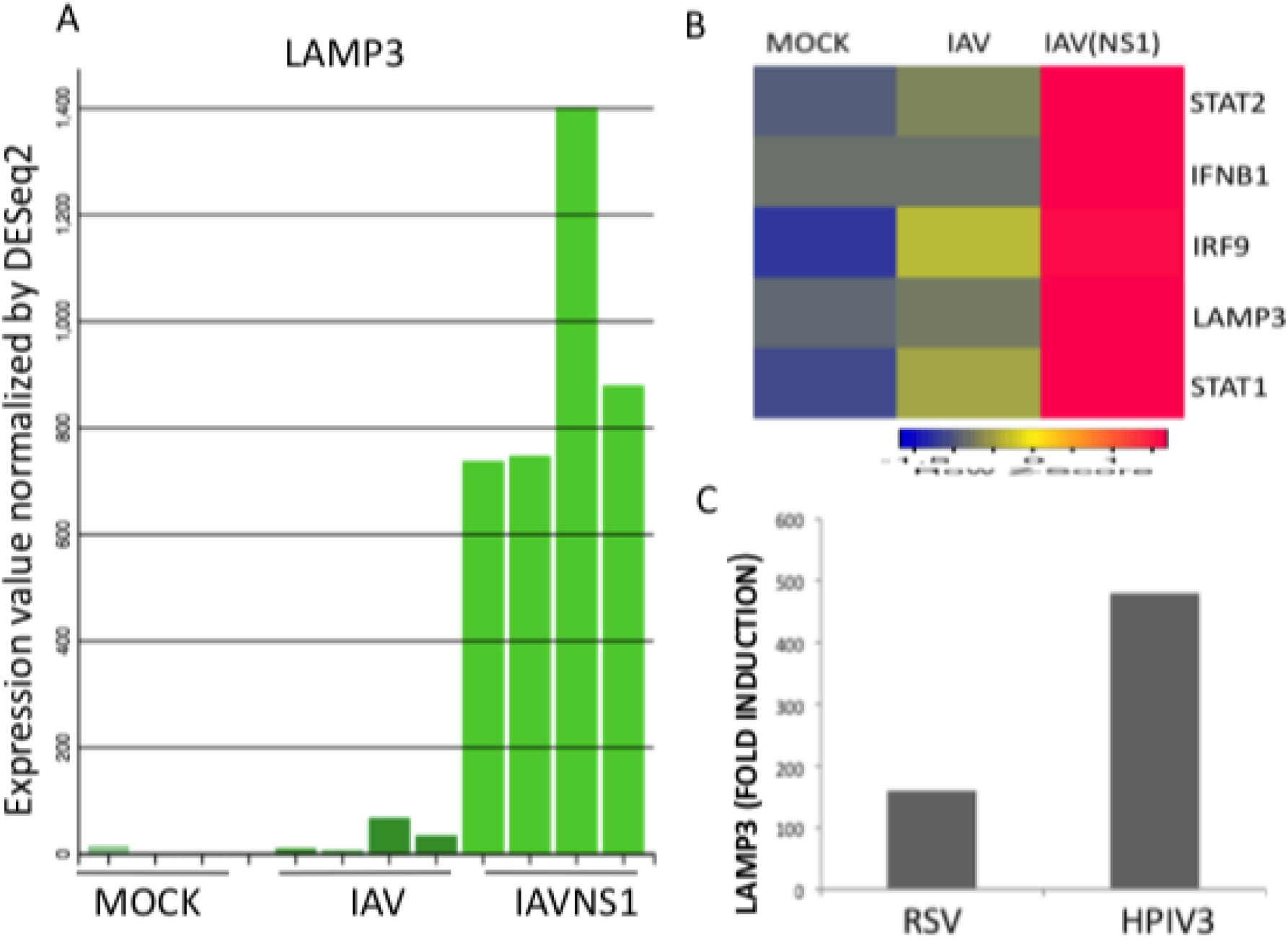
Regulation of LAMP3 mRNA levels by influenza viruses, respiratory syncytial virus (RSV), and the human parainfluenza virus (HPIV3). (A) Human NHBE1 bronchial epithelial cells were mock-infected or infected with influenza virus (IAV) or influenza virus lacking NS1 (IAVNS1) for 12 hours. Results from four samples for each condition were shown (B) Fold change in RNA expression levels of LAMP3, IFNB1 and ISGF3 transcription factor complex mRNA in NHBE cells after infection with wild type or NS1 deleted influenza A virus for 12 hours versus mock-infected controls. Average fold induction from four samples was shown (C) Human A549 lung cells were mock-infected or infected with RSV or HPIV3 for 24 hrs. LAMP3 mRNA levels were normalized by DESeq2.

### 3.6 Regulation of LAMP3 expression by Interferons

There are three distinct types of interferons-type I Interferon (IFN-α/β), type II interferon (IFN-γ), and type III interferon (IFN-λ).. Interrogation of microarray datasets revealed that IFN-β treatment induced LAMP3 expression in NHBE and human lung epithelial cells. Expression was induced by 4-6 hr and further enhanced by 24 hours (Figure 8A and 8B). Expression data in Interferon resource (Interferome) revealed that LAMP3 was regulated by type I and type II interferon but not type II interferon (data not shown). Transcriptional regulation by type 1 interferons is mediated by Stat, Stat2, and IRF9 trimeric complex binding to a regulatory element known as interferon-stimulated response element (ISRE) or Stat1 homodimer binding to another regulatory element known as gamma-activated sequence (GAS) located near transcription start site (TSS) of interferon-stimulated genes (45). Transcription factor binding site data in the promoter region of LAMP3 revealed that there were no ISRE or GAS elements within the 1.5 Kb upstream of the transcription start site. Interestingly, NF-kB and IRF-8 binding sites were detected. In contrast, consensus binding sites for Stat1 or Stat3 were detected immediately downstream of the transcription start site (Figure 8C). However, promoter analysis and chromatin immunoprecipitation experiments are required to confirm the role of interferon response elements and the binding of STAT proteins in LAMP3 gene regulation. Alternatively, novel regulatory elements and transcription factors may be involved in interferon regulation of LAMP3 gene (45). Bioinformatic analysis revealed that alterations in cell cycle and antiviral genes were responsible for a majority of cervical cancers and LAMP3 as a regulator of antiviral gene expression (46). In Hela cervical carcinoma cells, IFN-α treatment induced antiviral genes including STAT1, IRF7, EIF2AK2 (PKR), and LAMP3. Furthermore, knock-down of LAMP3 expression by Si-RNA prevented the induction of antiviral gene expression by IFN-α treatment in Hela cells (46).

**Figure 8.**
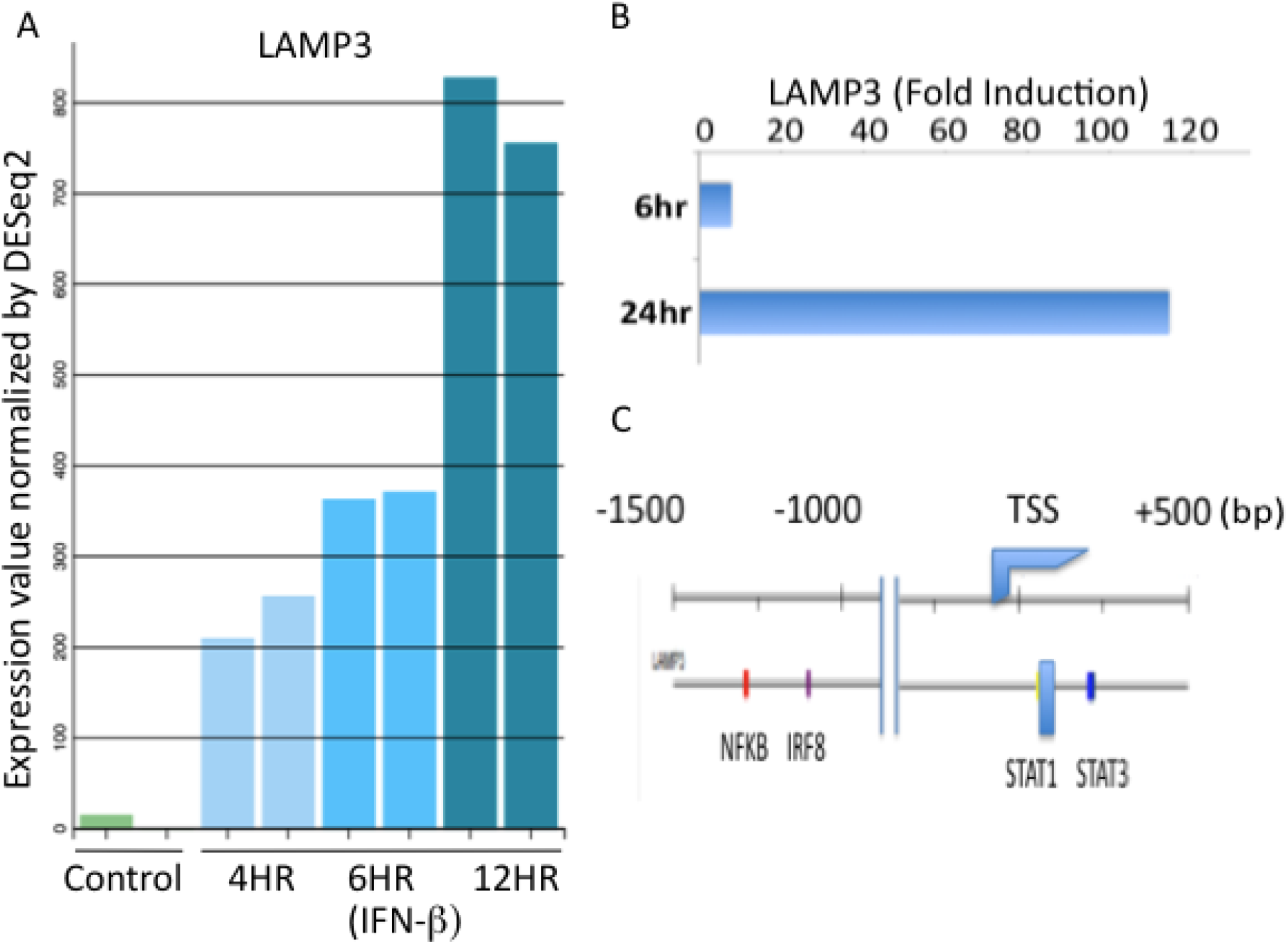
Regulation of LAMP3 mRNA levels by interferon-beta (IFN-β) treatment in human lung epithelial cell lines. (A) Regulation of LAMP3 mRNA levels by IFN-β treatment for 4, 6, or 12 hours in human NHBE cells (B) Regulation of LAMP3 mRNA levels by IFN-β treatment for 6 or 24 hours in human A549 cells (C) Schematic representation of the human LAMP3 gene promoter. The approximate location of the NF-kB, IRF8, STAT1 and STAT3 binding sites was shown. TSS represents the transcription start site. The map is not to scale.

### 3.7 Protein-protein interaction network of LAMP3

Signal transduction in mammalian cell systems is a complex process involving protein-protein interactions resulting in post-translational modifications and alteration of biological function (21,22). Protein interactions provide novel insights into the biological function of the proteins involved in the interactions. These interactions are regulated by factors such as location, proximity and are affinity-driven processes. Protein interactions of LAMP3 were visualized in the STRING database (Figure 9). These include interactions with a wide variety of CD family transmembrane glycoproteins involved in antigen presentation that are structurally related to major histocompatibility complex (MHC) proteins. Membrane channels and transporters are involved in transporting ions and solutes and play an important role in the absorption, distribution, and excretion processes. Several ATP-binding cassettes (ABC) containing proteins like ATP binding casette sub family A member 3 (ABCA3) and solute carrier transporters like SLC34A2 were highly expressed in type II cells (47). LAMP3 interacts with transporter protein ABCA3 involved in lipid and cholesterol metabolism which in turn interacts extensively with surfactant proteins. This network also includes the solute transport protein SLC34A2. Phosphate is a major constituent of surfactant and intake and extrusion of inorganic phosphate is determined by the SLC34A2, a sodium-phosphate cotransporter. LAMP3 protein interactions visualized in BIOGRID are extensive and encompass many other biological functions. LAMP3 protein interaction with janus kinase 1 (JAK1) tyrosine kinase involved in both type I and type II interferon signaling was reported in BIOGRID (data not shown). If this interaction is experimentally validated by different methods it might shed light on the mechanism by which LAMP3 regulates interferon signaling, constituting a feedback loop.

**Figure 9.**
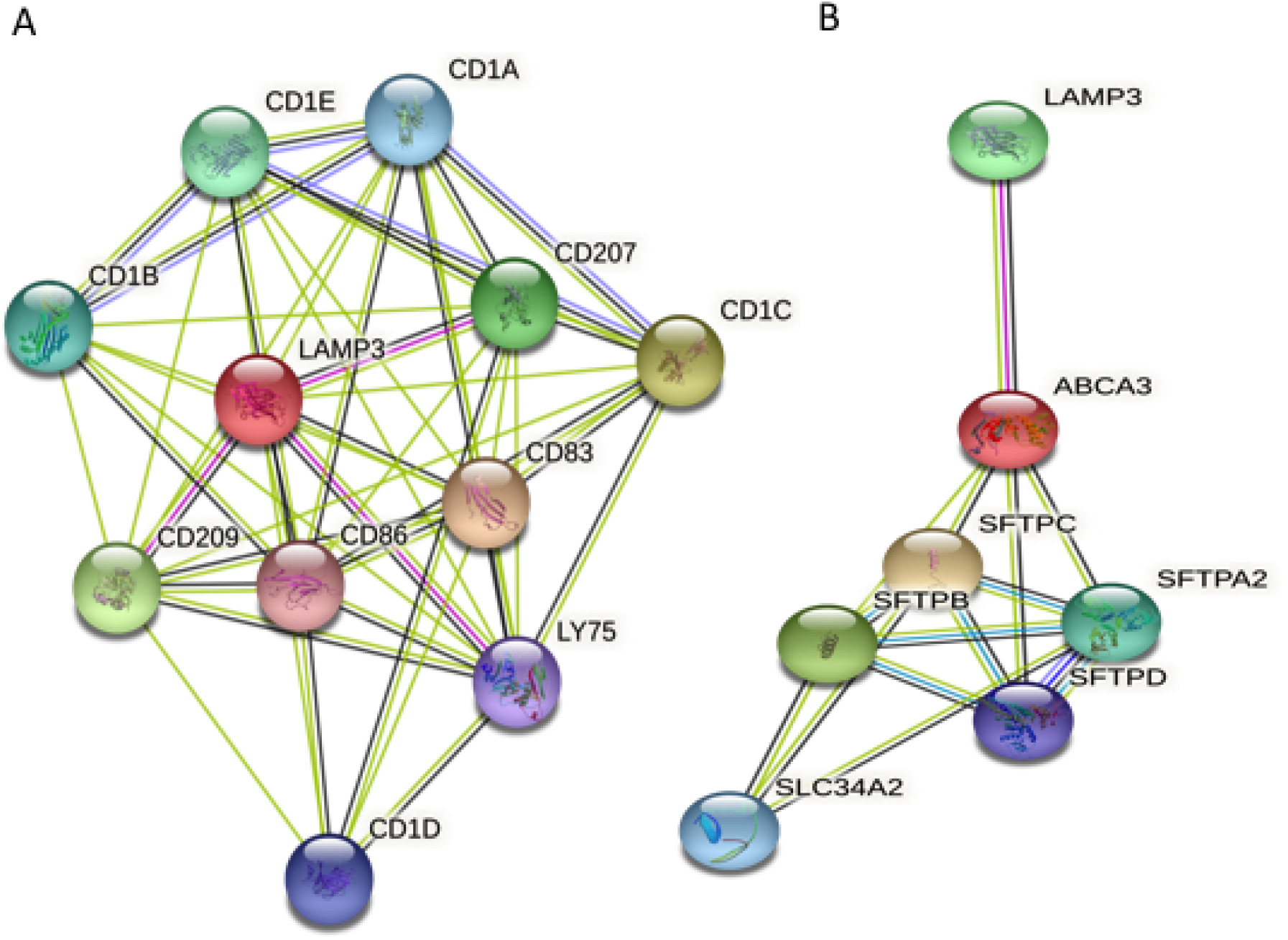
Protein-protein interactions of LAMP3 and other proteins in immune cells and lung type II cells were represented in the STRING database. Interacting proteins were represented by ovals and protein interactions by edges, respectively.

### 3.8 Regulation of lung type II transcriptome in Healthy and COVID-19 patients

Several cell culture models were developed to understand the role of SARS-CoV-2 replication and its viral proteins in reprogramming the lung type II transcriptome. These studies were done in ACE2 reconstituted A549 cells that were derived from human lung adenocarcinoma (48). These systems provided useful information on the direct effects of the virus replication on lung type II transcriptome. Lung injury in humans is mediated by the direct effects of the virus as well as the host immune responses. Mouse and other animal models were also limited in accurately reflecting transcriptome changes in human epithelial type II transcriptome. As an alternative approach, a list of human genes that correspond to mouse lung type II cell-specific genes as well as genes that represent major functional categories of type II cell such as transcription factors, metabolism, and redox regulation was interrogated in a microarray dataset of lung autopsies in healthy and COVID-19 patients (15,16).

Differential regulation of several membrane proteins including transporters and receptors were observed in healthy and COVID-19 patients. These include up-regulation of LAMP3, SLC34A2, and ATPase H+ transporting V1 subunit B2 (ATP6V1B2) and down-regulation of platelet derived growth factor C (PDGFC), chloride voltage-gated channel 6 (CLCN6), and FERM, Arh/RhoGEF and pleckstrin containing protein 1 (FARP1). Among the secreted surfactant proteins SFTPA1, SFTPA2, and SFTPB were up-regulated and SFTPC was down-regulated. In contrast, transcription factors such as PIN2 (TERF1) interacting telomerase inhibitor 1 (PINX1), bromodomain and WD repeat domain containing 1 (BRWD1), and activation transcription factor 6B (ATF6B) were down-regulated (Figure 10). Lung type II cells have a major role in carbohydrate and lipid metabolism and convert glycogen into phospholipids that are a major component of the surfactant proteins (8). C/EBP alpha (CEBPA), sterol regulatory-element binding protein (SREBP1), and signal transducers and activators of transcription 3 (STAT3) knockout mice have defects in lipid metabolism in lung type II cells (49–51). For example, conditional deletion of STAT3 in mice resulted in the decreased expression of ABCA3 and abnormalities in the formation of lamellar bodies, the intracellular organelles that store surfactant lipids (51). Genes that were enriched in lung type II cell-specialized functional categories such as lipid metabolism (LCN2, PLP1, and ALAS1), carbohydrate metabolism (LYZ, CHIT1, and CH3L1), redox-regulation (HMOX1, GLRX, FTH1, SOD2), and immune responses (LY6E, SLPI, and EMB) were up-regulated (Figure 11). The transcription factors of lung type II cells including signal transducer and activators of transcription 1 (STAT1), activator protein 1 (AP1), and nuclear factor kappaB (NF-kB) family members are activated by phosphorylation and play major roles in the influenza virus and inflammatory cytokine signaling pathways that mediated lung injury (52,53). Furthermore, induction of transcription factor at mRNA levels provides another regulatory step in cytokine signaling (46). Studies from knock-out mice have shown that transcription factors such as AP1 and NFE2L2 (nuclear factor, erythroid 2 like 2 or NRF2) play major roles in detoxification and redox regulation (54,55). Targeted deletion of AP1 in alveolar epithelial cells of mice resulted in enhanced lung inflammation and progressive emphysema in response to cigarette smoke (54). The NRF2 transcription factor binding to antioxidant/electrophile responsive element (ARE) and glutathione (GSH) regulates the expression of a large number of antioxidant enzymes (56). In the transcription factors category STAT1, JUND, and STAT3 were up-regulated and activator protein (AP1) family members such as FOS, JUN, and JUNB were downregulated (Figure 11). These results indicate dramatic changes in the lung type II transcriptome and functional reorganization. A summary of the results was presented in Table 1.

**Table 1.**
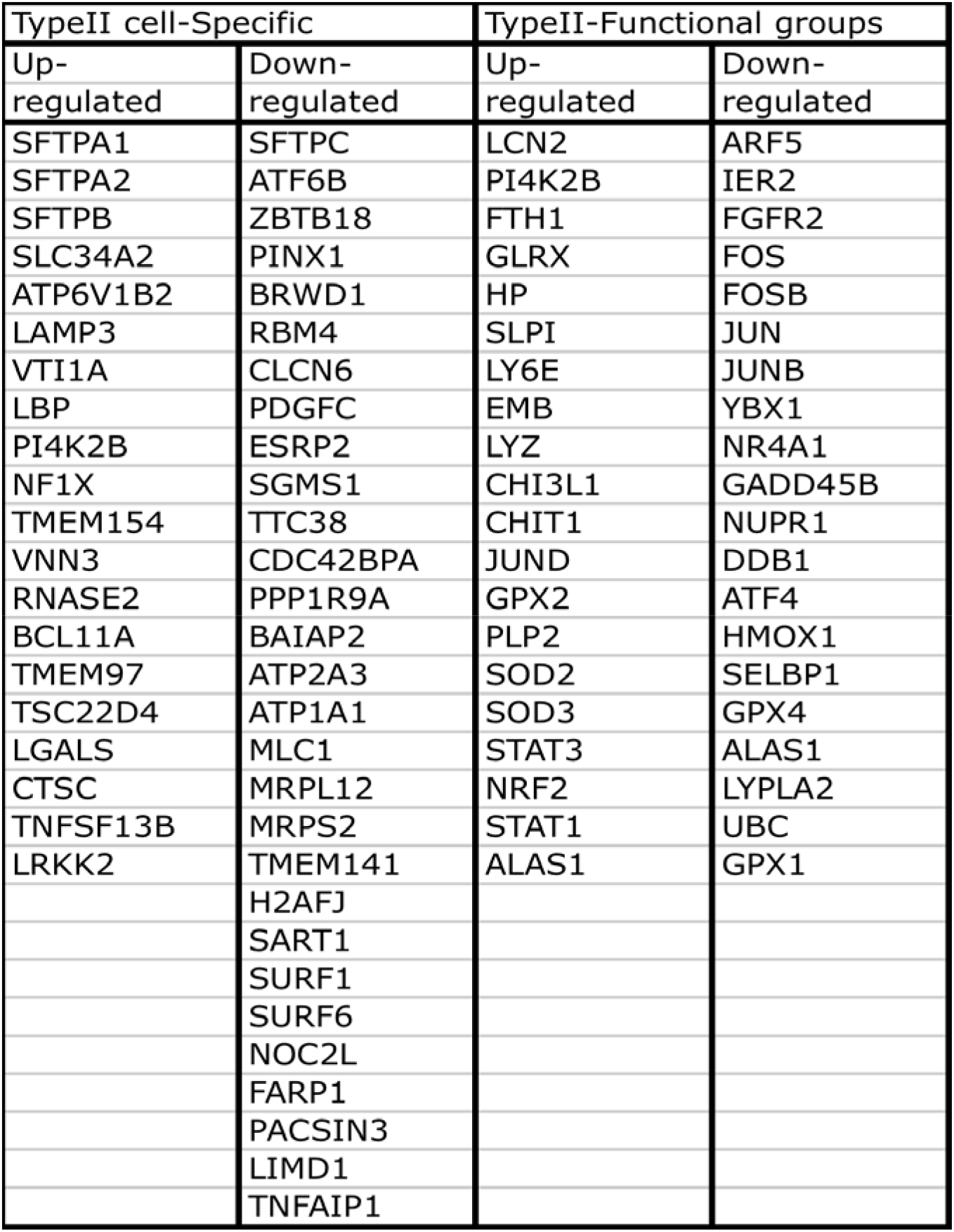
Regulation of lung epithelial type II-cell transcriptome in COVID-19 patients

**Figure 10.**
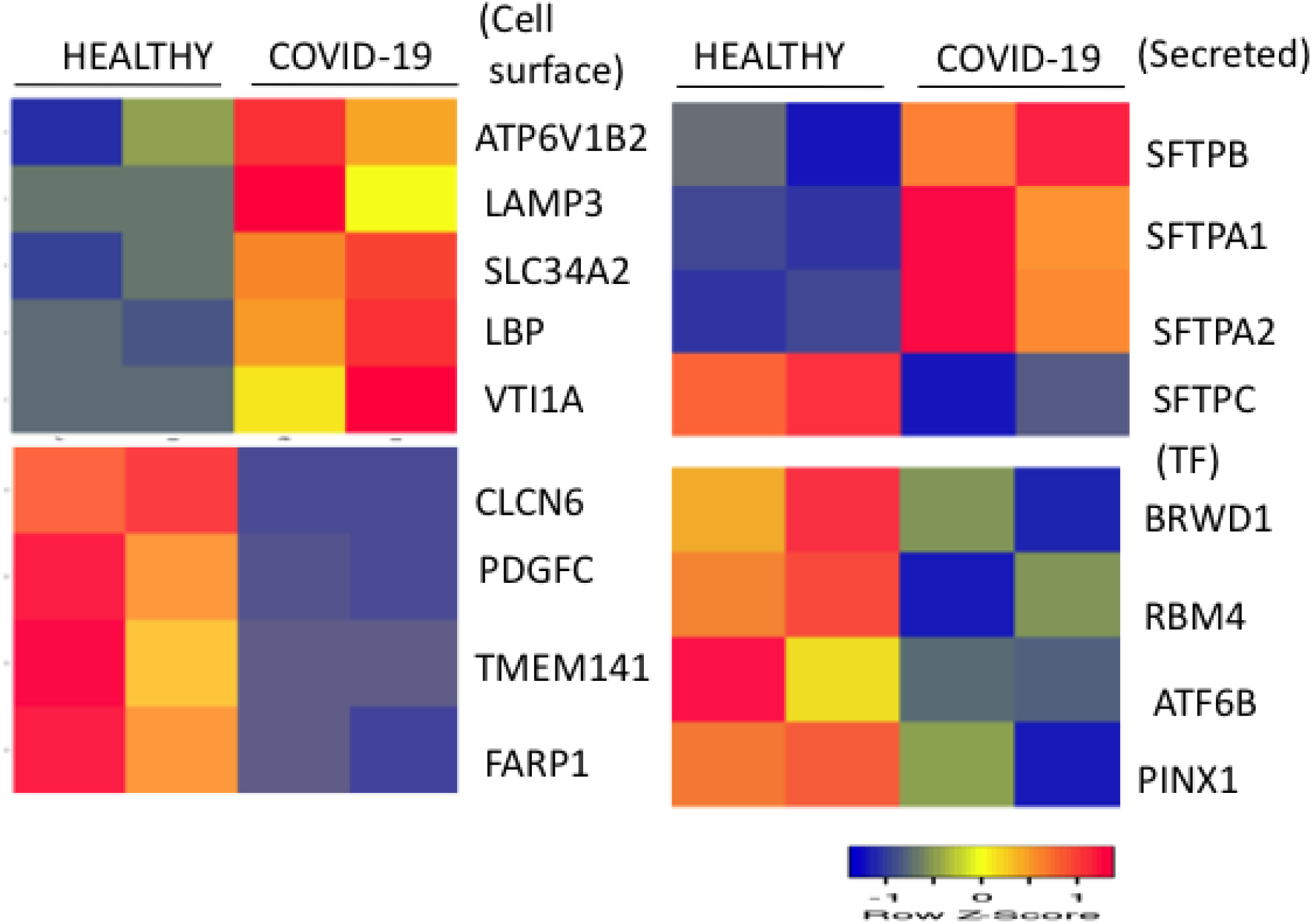
Regulation of lung type II-specific mRNA levels in healthy controls and COVID-19 patients. Expression levels of selected genes in healthy and COVID-19 lung samples were shown. Low and high expression in healthy and COVID-19 lung biopsies were represented by blue and red, respectively. RNA expression values were normalized by DEseq2 and shown as a heat map. Data from two samples for each condition were shown.

**Figure 11.**
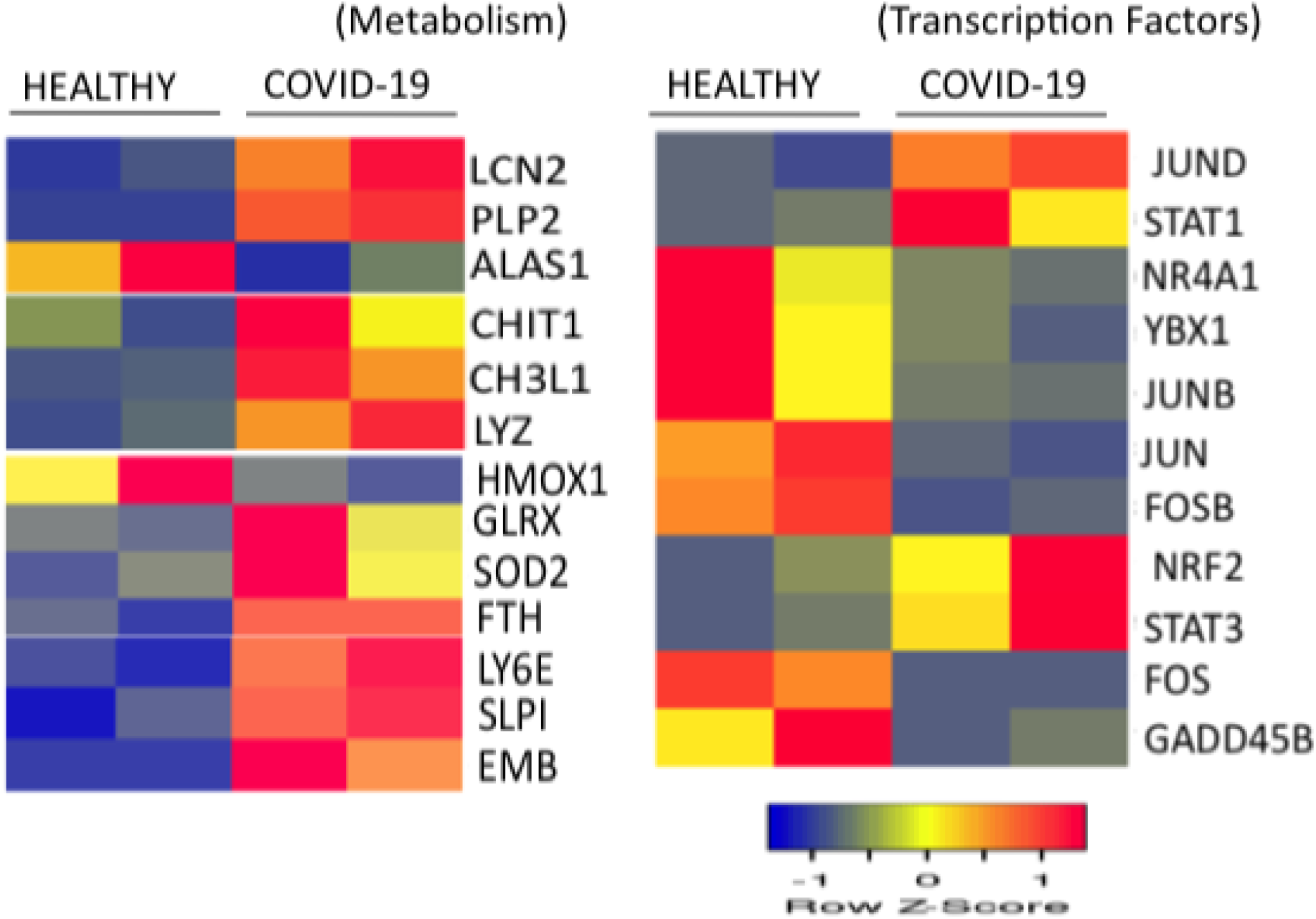
Regulated expression of the type II cell-enriched genes in the lung biopsies of healthy controls and COVID-19 patients. Expression levels of selected genes in healthy and COVID-19 lung samples were shown. Low and high expression levels were represented by blue and red, respectively. RNA expression values were normalized by DEseq2 and shown as a heat map. Data from two samples for each condition were shown.

Protein-protein interactions provide novel insights into the gene sub-networks in signal transduction pathways and biological functions in mammalian cells (19,20). Differentially regulated genes in the lungs of healthy and COVID-19 patients were investigated for potential protein interactions in the STRING database. Three sub-networks consisting of metabolism, oxidative stress, and transcription factors were represented in the network (Figure 12). Gene ontology terms (GO) associated with this interacting network include myeloid leukocyte activation, cellular oxidant detoxification, and diseases associated with surfactant metabolism (Figure 13A). Examination of the sub-network protein interactions revealed extensive interactions of carbohydrate and lipid metabolism (LYZ, CHIT11, and LCN2) with haptoglobin (HP) and secretory leukocyte peptidase inhibitor or SLPI (Figure 13B). Reactive oxygen species (ROS) including the superoxide, hydroxyl radicals and hydrogen peroxide generate oxidative stress resulting in toxic effects and cellular damage to macromolecules (57). Several antioxidant proteins including superoxide dismutase (SOD), glutathione peroxidase (GPX) Peroxiredoxin (PRX), glutaredoxin (GLRX), and catalase (CAT) are expressed in lung type II cells to minimize the oxidative damage in the lung (15). The redox system of glutathione and associated proteins are involved in regulating detoxification and antioxidant defense in lung type II cells. Its depletion in the lung has been associated with an increased risk of lung injury and disease (57,58). Protein interaction network analysis involved in redox regulation revealed that heme oxygenase 1 or HMOX1 plays a major role with connections to SOD3 and ferritin heavy chain 1 (FTH1) of antioxidant network and LCN2, 5’ aminolevulinate synthase 1 (ALAS1), and STAT3 of lipid metabolism network (Figure 13C). Consistent with these results reactive oxygen species and oxidative stress have been implicated in modulating lung type II cell metabolism and lung diseases (57,59,60).

**Figure 12.**
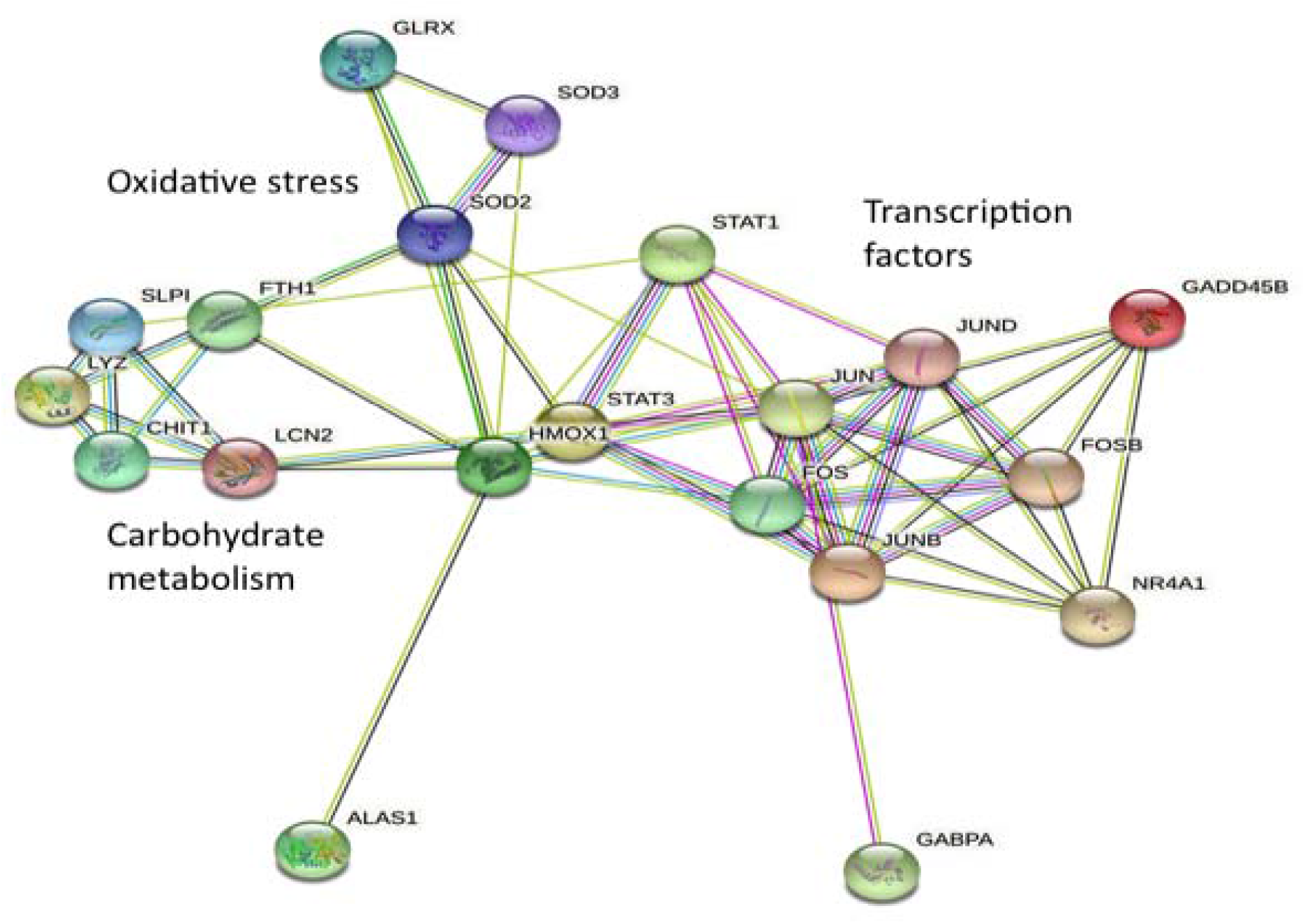
Protein-protein interactions and sub-networks of genes differentially regulated in the lungs of healthy and COVID-19 patients were represented in the STRING database. Transcription factor, oxidative stress, and carbohydrate metabolism sub-networks were indicated. Interacting proteins were represented by ovals and protein interactions by edges, respectively.

**Figure 13.**
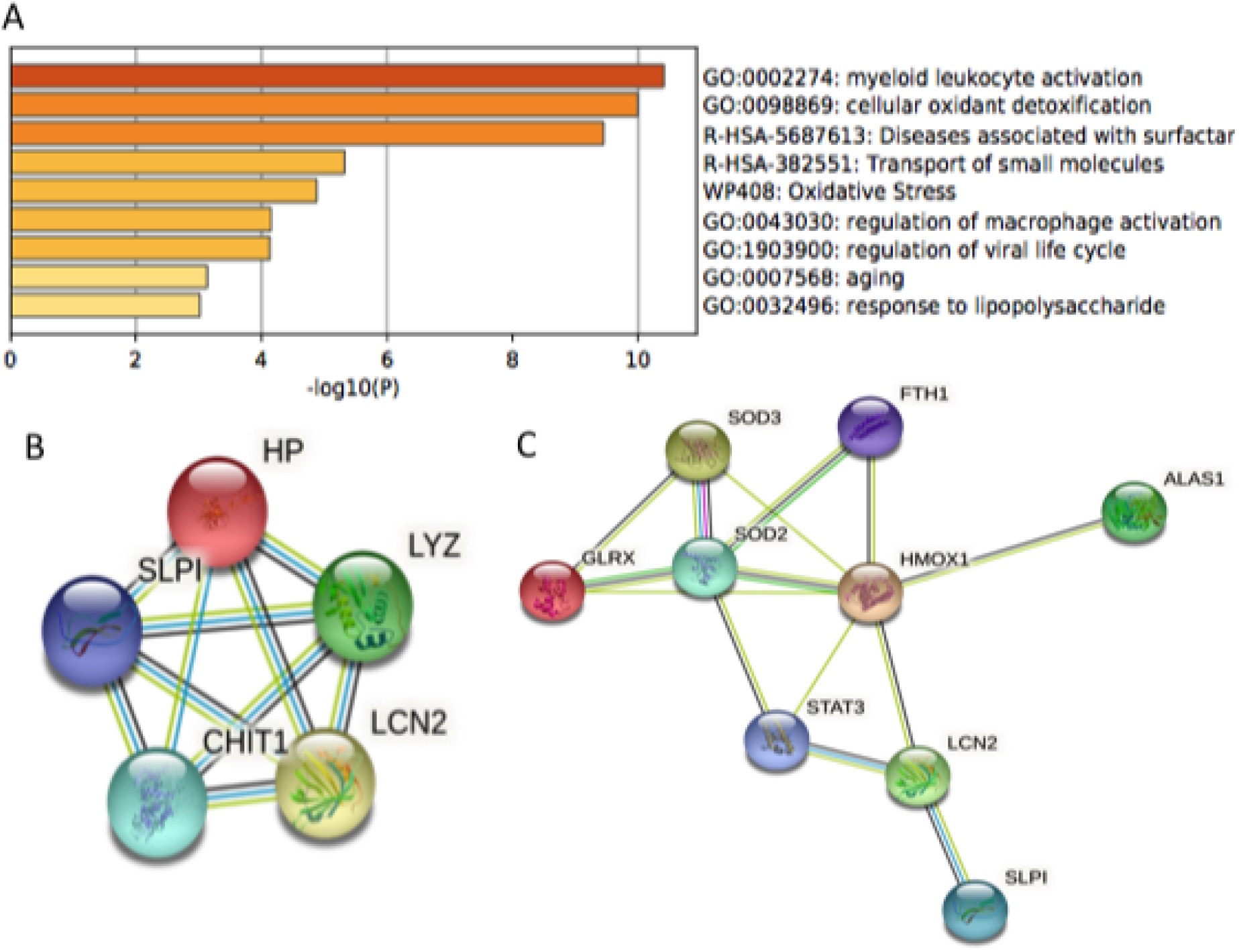
Gene ontology (GO) terms and protein interaction sub-networks of genes differentially regulated in the lungs of healthy and COVID-19 patients (A) Gene ontology (GO) terms associated with lung type II genes differentially regulated in healthy and COVID-19 patients were represented in Metascape database (B) Protein interactions of carbohydrate metabolism and oxidative stress sub-networks as represented in the STRING database

## 4. CONCLUSION

Activation of transcription factor sub-networks in type I interferon signaling plays an important role in innate immunity to respiratory viruses (46). Transcription factors of STAT, NF-KB, AP1, and ATF family are activated in lung epithelial cells in innate and adaptive immunity leading to the production of chemokines and cytokines that in turn recruit inflammatory cells such as macrophages, neutrophils, and others (39,40). Profiling lung epithelial type II-specific genes led to the identification of lysosomal-associated membrane protein 3 (LAMP3) and its role in vesicular transport. LAMP3 expression was induced in human lung epithelial cells by several respiratory viruses including SARS-CoV-2, influenza A, RSV, and HPIV3. LAMP3 expression was induced by type I interferon in human lung epithelial cell lines. Hypoxia regulated LAMP3 via PKR-like endoplasmic reticulum kinase (PERK), Activation transcription factor 4 (ATF4) and unfolded protein response (UPR) pathway in head and neck squamous cell carcinoma (61). Furthermore, promoter analysis and chromatin immunoprecipitation assay (CHIP) demonstrated that ATF4 regulated LAMP3 expression (62). These studies suggest that LAMP3 may be involved in immune evasion in response to pathogens and cancer. Microbiota associated with cervical cancers includes *Prevotella bivia* and co-culture of *P.Bevia* with cervical carcinoma cells induced STAT1 and LAMP3 (63). Metagenomic data in COVID 19 patients revealed that *Prevotella sp* was a common opportunistic secondary infection in the lungs (64). Furthermore, knock-down of LAMP3 in human cervical carcinoma cells down-regulated interferon-stimulated gene expression suggesting a critical role in interferon signaling (46). Location in lysosomes and endosomes, as well as regulation by pathogenic bacteria and respiratory viruses, suggests that LAMP3 may have an important role in inter-organellar regulation of innate immunity. Targeting LAMP3 and specific components of vesicular transport may have applications in human health and disease.

